# Endogenous retrotransposon activity supports osteocyte function and links antiretroviral therapy to bone loss

**DOI:** 10.64898/2025.12.27.696700

**Authors:** Arianna Mangiavacchi, Sjur Reppe, Gabriele Morelli, Marco D’Onghia, Kaare M. Gautvik, Valerio Orlando

**Affiliations:** King Abdullah University of Science and Technology (KAUST), Biological Environmental Science and Engineering Division, Thuwal, 23500-6900, Kingdom of Saudi Arabia; Oslo University Hospital, Department of Medical Biochemistry, Oslo, Norway; Lovisenberg Diaconal Hospital, Unger-Vetlesen Institute, Oslo, Norway; Oslo University Hospital, Department of Plastic and Reconstructive Surgery, Oslo, Norway

**Author notes:** Corresponding authors: Arianna Mangiavacchi, Valerio Orlando.

## Abstract

Retrotansposable elements such as LINE-1 and HERV-K encode endogenous reverse transcriptases (RTs) with emerging roles in human biology. People living with HIV, particularly those receiving nucleoside reverse transcriptase inhibitor (NRTI)-based therapy, have increased risk of low bone mineral density (BMD), but the underlying mechanisms remain unclear. Here we show that endogenous RT activity, estimated by L1-Ta and HERV-K DNA content, is enriched in human bone and markedly reduced in osteoporosis, where it correlates with BMD and osteocyte gene expression. *In vitro*, nucleoside reverse transcriptase inhibitors (NRTIs), widely used as antiretroviral drugs, impair osteocyte endocrine function, disrupting vitamin D_3_–induced FGF23 production and parathyroid hormone– mediated regulation of SOST. These findings identify endogenous RT activity as a regulator of osteocyte function and bone homeostasis, and suggest that its pharmacological inhibition may contribute to antiretroviral therapy–associated bone loss.

## Introduction

People living with human immunodeficiency virus (HIV) experience a substantially increased risk of low bone mineral density (BMD), skeletal fragility, and fragility fractures across all age groups, particularly when receiving nucleoside reverse transcriptase inhibitor (NRTI)–based antiretroviral therapy (ART) ^1 2 3 4 5 6 7 8^. Significant effects of NRTIs on bone metabolism have been reported *in vitro* ^9 10^ and *in vivo* using a macaque model of HIV infection, in which NRTI monotherapy resulted primarily in defective mineralization of newly formed bone ^11 12 13^. Although multifactorial mechanisms have been proposed, including direct antiretroviral toxicity, the etiology of ART-associated secondary osteoporosis remains incompletely understood. NRTIs, including Tenofovir (TDF), Abacavir (ABC), and Lamivudine (3TC) inhibit HIV reverse transcriptase by acting as nucleotide analogues of ATP, GTP, or CTP, respectively, that are incorporated into the nascent viral DNA strand, thereby terminating further DNA synthesis ^14 15^. However, previous studies have demonstrated the ability of several NRTIs to also target endogenous reverse transcriptases (RT), particularly those derived from LINE-1 (long interspersed nuclear element-1) and ERV-K (human endogenous retrovirus K) retrotransposons, two major contributors to endogenous reverse transcription activity in mammalian cells ^16 17 18 19 20 21^. Transposable elements (TEs), which constitute 50% of the mammalian genome,^22^ are increasingly appreciated as transcriptionally active genomic components with physiological functions extending beyond their classical role as mobile genetic elements.^23^ While historically viewed as genomic parasites, TE are now recognized as contributors to cellular homeostasis, stress responses, and tissue-specific functions.^24 25 26 27 28^ We recently demonstrated that TE transcription, particularly from LINE and HERV families, is sensibly reduced in osteoporotic bone and that exogenously provided TE RNAs support anabolic bone responses and potentially microfracture repair via a paracrine mechanism, highlighting a functional role for these elements in skeletal biology.^29^ Notably, LINE-1 and HERV-K are unique in encoding functional RTs, enabling endogenous reverse transcription of their RNAs into cDNA in somatic cells.^21 30 31^ NRTI modulation of endogenous RT activity has already been shown to impact the phenotypes of several murine models, included attenuation of retroelement-driven inflammation and joint damage in arthritis ^32^, rescue of learning deficits in mice with Down syndrome ^33^, amelioration of the aging phenotyope in Sirt6-/-mice,^34^ and inhibition of vascular calcification.^35^

Given the high prevalence of ART-associated osteoporosis and the known inhibitory effects of NRTIs on endogenous RTs, we hypothesized that LINE-1 and HERV-K–derived RT activity contributes to bone homeostasis and that its inhibition may lead to reduced bone integrity and fragility. We quantified L1-Ta and HERV-K DNA content as a proxy for endogenous RT activity in human bone biopsies and examined its relationship with BMD, aging, and osteocyte gene expression. Furthermore, we used NRTIs in a validated osteoblast-to-osteocyte differentiation model to assess the functional impact of RT activity on osteocyte maturation and endocrine responsiveness. Together, our findings reveal endogenous RT activity as a novel component of osteocyte biology and identify its inhibition as a potential drawback effect affecting bone integrity in clinical practice, particularly in long-term exposure to NRTIs.

## Results

### L1-Ta and ERV-K DNA content is enriched in human bone and shows variation with individual age

LINE-1 (L1-Ta) and HERV-K (HML-2) elements encode functional RT and constitute the major source of endogenous RT activity in human cells ^21 31^. Reverse transcription of TE-derived RNA produces complementary DNA (cDNA) that can persist as extrachromosomal intermediates or integrate into the genome, thereby increasing TE DNA content, which can serve as a proxy for endogenous RT activity. Since endogenous RTs, particularly ORF2p, are typically present at levels too low to be reliably detected and quantified using standard protein-based methods,^36 37 38 39 40^ the most sensitive and reliable approach, albeit indirect, to assess endogenous RTs activity is to measure the DNA/cDNA content of their targets. Therefore, we quantified L1-Ta and HERV-K DNA content in trans-iliac bone biopsies from 84 demographically matched postmenopausal women in comparison with adjacent skeletal muscle and peripheral blood mononuclear cells (PBMCs) from the same donors. The donors were a homogeneous group with regard to diet, nutritional supplements, and lifestyle factors, including physical engagement ^41^. Compared to muscle tissue and PBMCs, the DNA content of L1-Ta and HERV-K was significantly enriched in bone, indicating higher endogenous RT activity in this tissue (Fig. 1A, B). As expected from their genomic abundance, L1-Ta copies were more numerous than HERV-K (Supplementary Fig. S1). Notably, correlation analysis showed a progressive decline of both L1-Ta and HERV-K content with age in bone, but not in muscle or blood, suggesting that age-related decrease in RT activity is bone-specific (Fig. 1 C, D; Supplementary Fig. S2). Similarly, BMD decreased with age at the total body, hip and lumbar spine (Fig. 1E), supporting an association between reduced endogenous RT activity and age-related bone loss.

**Fig. 1.**
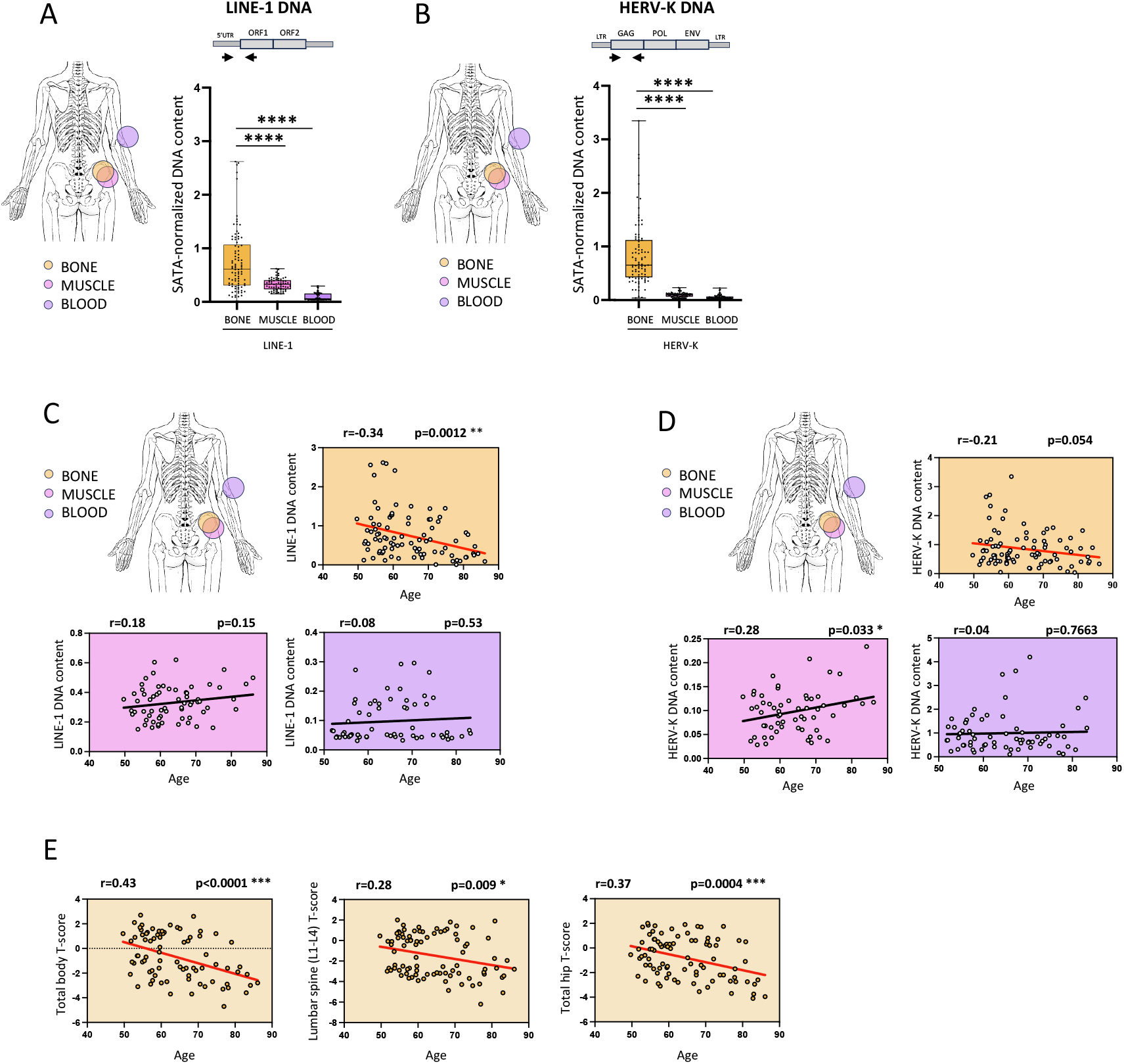
Tissue-specific L1-Ta and HERV-K DNA content and correlation with individual age. A) L1-Ta DNA content in bone, muscle and blood biopsies. Values are plotted over the average of bone. Mean (bone= 1; muscle=0.305; blood=0.094). Statistically significant difference between mean ± SEM (bone vs muscle: -0.6946 ± 0.1241; bone vs. blood: -0.906 ± 0.12). B) HERV-K DNA content in bone, muscle and blood biopsies. Values are plotted over the average of bone. Mean (bone= 1; muscle=0.115; blood=0.063). Statistically significant difference between mean ± SEM (bone vs muscle: - 0.7496 ± 0.07767; bone vs. blood: -0.7936 ± 0.07315). C) Correlation analysis between L1-Ta DNA content and individual age in bone, muscle and blood biopsies. Correlation coefficient (r) is shown in each plot. D) Correlation analysis between HERV-K DNA content and individual age in bone, muscle and blood biopsies. Correlation coefficient (r) is shown in each plot. E) Correlation analysis between individual age and total body, spine and hip BMD t-score. Correlation coefficient (r) is shown in each plot.

### L1-Ta and HERV-K DNA content is reduced in osteoporotic bone and correlates with individual BMD

We next stratified participants into healthy (n=46, BMD T-score >-1), osteopenic (n=20, − 2.5 < BMD T-score ≤ − 1), and osteoporotic (OP, n=18, BMD T-score ≤ − 2.5) groups based on BMD T-scores and fracture history. Serum biomarkers confirmed active bone resorption in the OP group (Supplementary Table S1). L1-Ta DNA content was significantly reduced in osteoporotic bone compared with healthy controls (Fig. 2A). A similar reduction was observed for HERV-K (Fig. 2B). Compared to healthy bone, L1-Ta and HERV-K DNA content was markedly lower in PBMCs as well as in muscle, but most importantly, no differences were observed between healthy and OP groups within these tissues (Fig. 2C, D). Thus, the results show that a quantitative variation in L1-Ta and HERV-K DNA content is detected only in bone between osteoporotic patients and healthy donors, and not in two other mesoderm-derived tissues that are unaffected by osteoporosis. Therefore, alterations in endogenous RT dynamics in osteoporosis appear to be bone-specific and not confined to one skeletal site. Furthermore, we found that L1-Ta and HERV-K DNA content in bone correlates positively with BMD at total body, hip and lumbar spine (Fig. 3A,B), while significant correlation was not observed with individual parameters not strictly related to skeletal metabolism as body weight and serum vitamin D content (Supplementary Fig. S3). Expression of osteoblastic and osteocyte genes in the bone of healthy and OP participants was then quantified (Fig. 3C; Supplementary Table S2). Osterix (*SP7*) and Runt-related transcription factor 2 (*RUNX2)*, encoding essential transcription factors needed for osteoblast differentiation, were reduced in patients as well as Osteopontin (*SPP1*), commonly expressed in mature osteoblasts and osteocyte. However, the most pronounced reduction involved the osteocyte-specific markers Sclerostin (*SOST*) and Matrix Extracellular Phosphoglycoprotein (*MEPE)* (Fig. 3C; Supplementary Table S2). These results reveal a functional insufficiency in the mature osteoblasts/osteocytes anabolic activity as an important cause to bone fragility in this group of patients. L1-Ta and HERV-K DNA content correlated strongly with osteocyte-specific gene expression, particularly SOST and MEPE (Fig. 3D; Supplementary Fig. S4 A–B), while correlations with osteoblast genes did not reach significance (Fig. 3D; Supplementary Fig. S4 A–B). Together, these results link reduced mineral density, and particularly reduced osteocyte activity, to decreased RT dynamics in osteoporotic women compared to healthy women as measured in their bone biopsies.

**Fig. 2.**
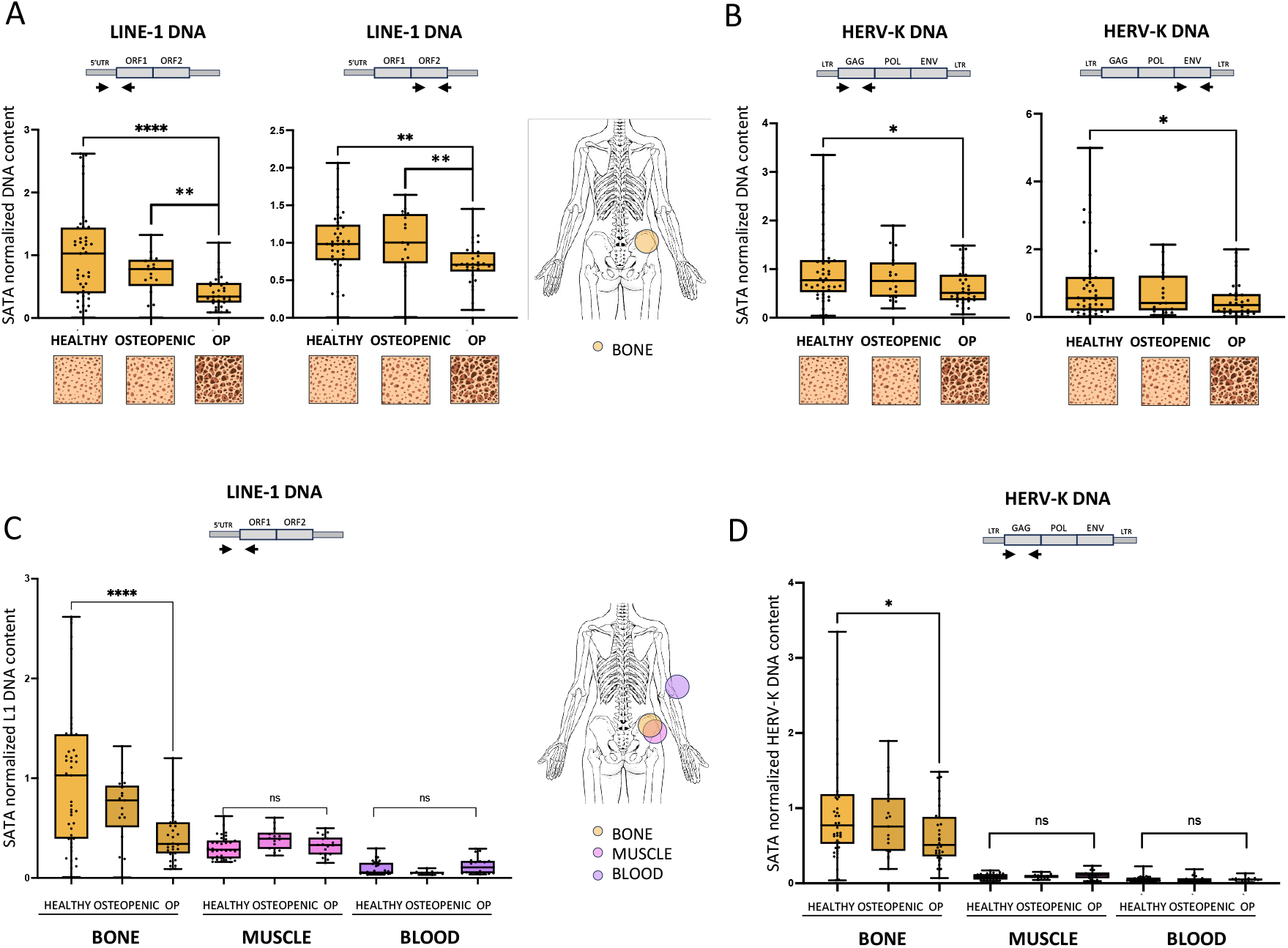
Tissue specific L1-Ta and HERV-K DNA content in healthy donors, osteopenic and osteoporotic patients. A) L1-Ta DNA content in bone biopsies from healthy, osteopenic, and osteoporotic patients. Values are plotted over the average of healthy. Left panel: Mean (healthy= 1; osteopenic= 0.694; op= 0.417). Statistically significant difference between mean ± SEM (healthy vs op= -0.5829 ± 0.1365; osteopenic vs op= -0.2767 ± 0.08703). Right panel: Mean (healthy= 1; osteopenic= 1.023; op= 0.7170). Statistically significant difference between mean ± SEM (healthy vs op= -0.2830 ± 0.08894; osteopenic vs op= -0.3060 ± 0.1000). B) HERV-K DNA content in bone biopsies from healthy, osteopenic, and osteoporotic patients. Values are plotted over the average of healthy. Left panel: Mean (healthy= 1; osteopenic= 0.833; op= 0.64). Statistically significant difference between mean ± SEM (healthy vs op= -0.3601 ± 0.1493. Right panel: Mean (healthy= 1; osteopenic= 0.7208; op= 0.53). Statistically significant difference between mean ± SEM (healthy vs op= -0.4700 ± 0.2255). C) L1-Ta DNA content in bone, muscle and blood biopsies of healthy, osteopenic and osteoporotic patients. Values are plotted over the average of healthy bone. Mean bone (healthy= 1; osteopenic= 0.694; op= 0.417); mean muscle (healthy= 0.305; osteopenic= 0.391; op= 0.324); mean blood (healthy= 0.094; osteopenic= 0.055; op= 0.125). D) HERV-K DNA content in bone, muscle and blood biopsies of healthy, osteopenic and osteoporotic patients. Values are plotted over the average of healthy bone. Mean bone (healthy= 1; osteopenic= 0.833; op= 0.64); mean muscle (healthy= 0.089; osteopenic= 0.1; op= 0.113); mean blood (healthy= 0.054; osteopenic= 0.05; op= 0.056).

**Fig. 3.**
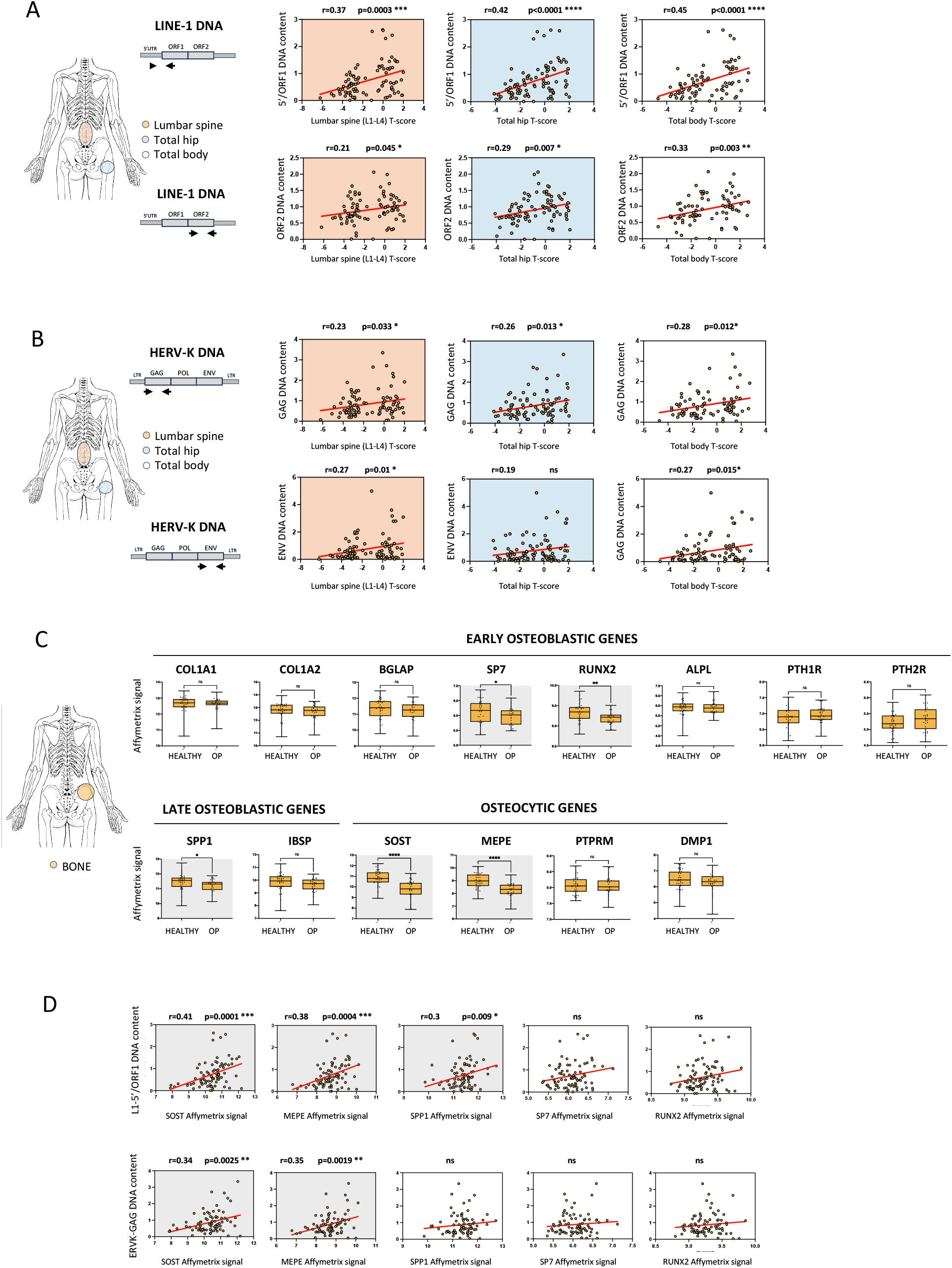
Correlation analysis between L1-Ta and HERV-K DNA content, bone mineral density (BMD) and expression of osteocytic genes in trans-iliac bone biopsies. A) Correlation between L1-Ta DNA content and spine, hip, and total body BMD T-score. Correlation coefficient (r) is shown in each plot. B) Correlation analysis between HERV-K DNA content and spine, hip and total body BMD T-score. Correlation coefficient (r) is shown in each plot. C) Microarray expression of early osteoblastic genes, late osteoblastic genes and osteocytic genes in bone biopsies from healthy women and osteoporotic patients. Mean signal levels and difference between mean ± SEM are listed in Table S2. D) Correlation analysis between DNA content of L1-Ta and HERV-K and the expression of genes downregulated in the biopsies of osteoporotic patients compared to healthy. Correlation coefficient (r) is shown in each plot.

### NRTIs alter osteocyte maturation and impair hormone responsiveness

Our analysis of human bone biopsies suggested a correlation between endogenous RT dynamics, osteocyte activity and bone homeostasis. Therefore, we tested whether pharmacological RT inhibition affected osteocyte maturation/function using Immortomouse/Dmp1-GFP-SW3 (IDG-SW3) cells, a validated model of osteoblast-to-osteocyte transition ^42^. Consistent with previous observations ^42^, when cultured under osteogenic conditions for 21 days, IDG-SW3 cells produced Dmp1-GFP, a marker of osteocytic differentiation ^43^ (Fig. 4A). They also deposited a highly mineralized extracellular matrix (Fig. 4A) and upregulated early (DMP1, MEPE, and PHEX) (Fig. 4B) and late (SOST and FGF23) osteocyte markers (Fig. 4C). Cells treated with the NRTIs TDF, ABC, or Lamivudine (3TC) during differentiation exhibited a selective reduction in FGF23 expression with 3TC and ABC, whereas TDF showed no significant effect. ABC increased DMP1 and SOST expression (Fig. 4D). Mineralization was slightly reduced by ABC and 3TC (Fig. 4E). When differentiated into late osteocytes, IDG-SW3 cells expressed SOST and FGF23, a critical regulator of phosphate homeostasis secreted by osteocytes and upregulated by 1,25-dihydroxyvitamin D_3_ and dietary phosphate intake ^44 45^. In mature osteocytes, 1,25-dihydroxyvitamin D_3_ induced a ∼15-fold increase in FGF23 expression in control cells (Fig. 4F). However, NRTI-treated cells exhibited blunted responses: TDF and 3TC induced only ∼6-fold increases, whereas ABC completely blocked FGF23 induction (Fig. 4G).

**Fig. 4.**
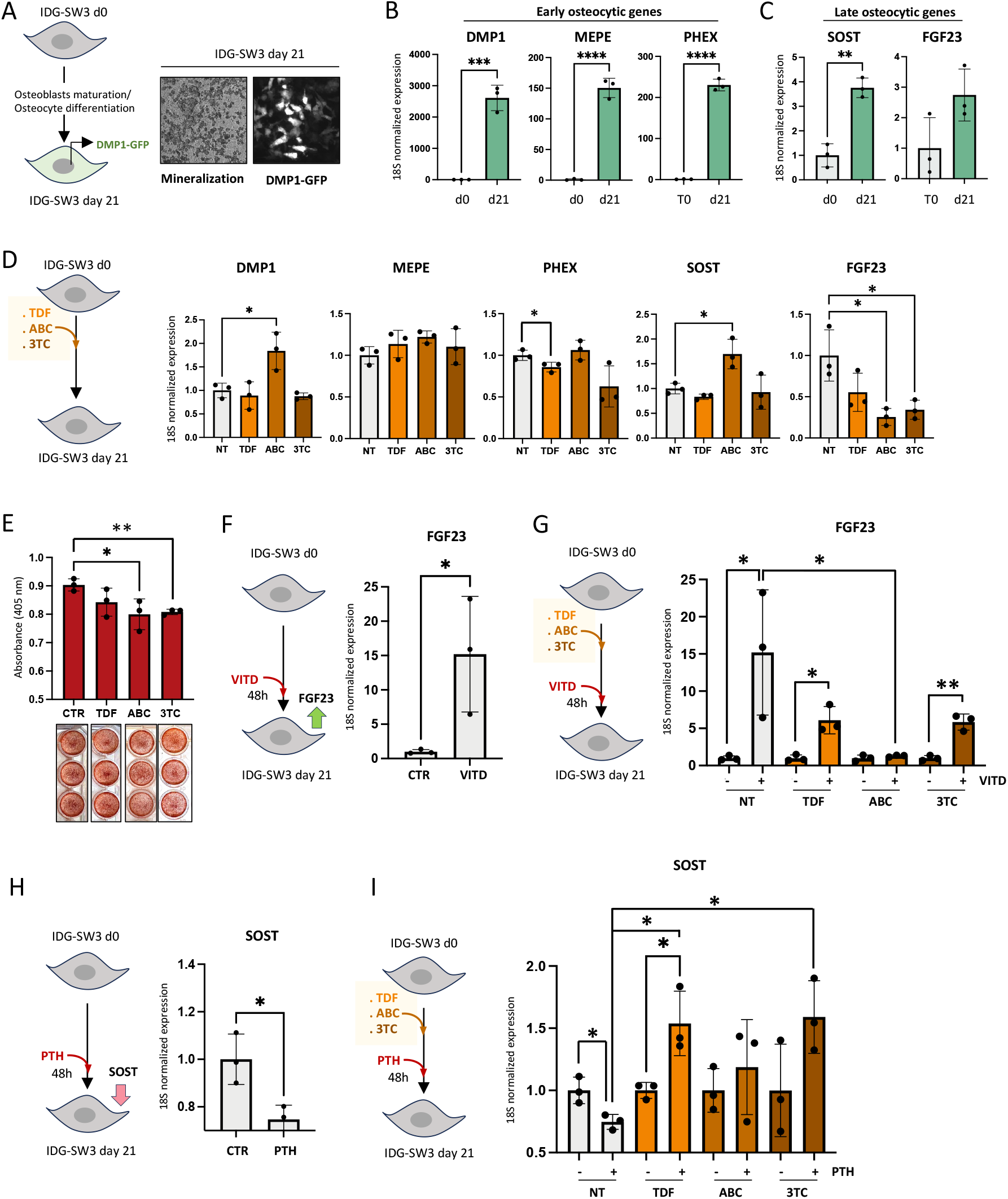
NRTI-treatement in IDG-SW3 osteocytes. A) Schematic representation of the experiment (left panel), mineralization and fluorescent signal of DMP1-GFP protein (right panel) in differentiated IDG-SW3 osteocytes. B) qPCR expression analysis of early osteocytic genes in differentiated (d21) IDG-SW3 osteocytes. Values are plotted over the average of undifferentiated cells (mean T0=1). Mean (DMP1= 2611; MEPE= 150.5; PHEX= 230.3) C) qPCR Expression analysis of late osteocytic genes in differentiated (d21) IDG-SW3 osteocytes. Values are plotted over the average of undifferentiated cells (mean T0=1). Mean (SOST= 3.757; FGF23= 2.744). D) Schematic representation of the experiments (left panel). qPCR expression analysis of early and late osteocytic genes in NRTIs-treated differentiated IDG-SW3 osteocytes. Values are plotted over the average of untreated cells (mean NT=1). Mean DMP1 (TDF= 0.89; ABC= 1.84; 3TC= 0.877). Mean MEPE (TDF= 1.136; ABC= 1.22; 3TC= 1.104). Mean PHEX (TDF= 0.859; ABC= 1.063; 3TC= 0.627). Mean SOST (TDF= 0.835; ABC= 1.697; 3TC= 0.928). Mean FGF23 (TDF= 0.554; ABC= 0.255; 3TC= 0.342). Tenofovir; ABC= Abacavir, 3TC = Lamivudine. E) Quantification (upper panel) of Alizarin red staining (lower panel) in control (NT) and NRTIs-treated differentiated IDG-SW3 osteocytes. Mean (CTR= 0.903; TDF= 0.842; ABC= 0.8; 3TC= 0.81). (F) Schematic representation of the experiment (left panel) and qPCR expression analysis of FGF23 in VITD-treated IDG-SW3 differentiated osteocytes (right panel). Values are plotted over the average of untreated cells (CTR). Mean (CTR=1; VITD = 2.744). G) Schematic representation of the experiments (left panel). qPCR expression analysis of FGF23 in untreated (NT) or NRTIs-treated differentiated IDG-SW3 osteocytes with (+) or without (-) VITD stimulation. Values are plotted over the average of CTR (-) groups. Mean (NT= 15.20; TDF= 6.101; ABC= 1.29; 3TC= 5.845). TDF= Tenofovir; ABC= Abacavir, 3TC = Lamivudine. H) Schematic representation of the experiment (left panel) and qPCR expression analysis of SOST in PTH-treated IDG-SW3 differentiated osteocytes (right panel). Values are plotted over the average of untreated cells (CTR). Mean (CTR=1; PTH = 0.747). I) Schematic representation of the experiments (left panel). qPCR expression analysis of SOST in untreated (NT) or NRTIs-treated differentiated IDG-SW3 osteocytes with (+) or without (-) PTH stimulation. Values are plotted over the average of CTR (-) groups. Mean (NT= 0.747; TDF= 1.538; ABC= 1.187; 3TC= 1.590). TDF= Tenofovir; ABC= Abacavir, 3TC = Lamivudine.

Previous studies have shown that SOST may play a role in mediating PTH action in bone ^46 47 48^. A single injection of PTH to mice *in vivo* was shown to reduce SOST expression and constitutive activation of the PTH receptor in osteocytes increased bone mass and reduced SOST expression ^49 50^. As shown in Figure 4H-I, the PTH-mediated suppression of SOST expression was preserved in control cells but disrupted in all NRTI-treated groups. TDF and 3TC increased SOST expression following PTH exposure, while ABC completely abolished the response.

Thus, while NRTIs had minimal effects on mineralization or early differentiation, they impaired key osteocyte endocrine functions, particularly the regulation of FGF23 and SOST by 1,25-dihydroxyvitamin D_3_ and PTH.

## Discussion

This study identifies endogenous reverse transcriptase (RT) activity derived from transposable elements as a previously unrecognized regulator of osteocyte function and bone homeostasis.

Osteocytes are master regulators of skeletal remodeling, integrating mechanical, hormonal, and metabolic signals to coordinate bone formation and resorption. Their ability to respond dynamically to mechanical loading and systemic cues requires sustained transcriptional plasticity and robust signaling networks. ^51 52 53^

Our findings suggest that TE-derived RT activity contributes to this regulatory capacity, extending previous observations that TE transcription supports anabolic bone responses and microfracture repair.^29^ The strong association between L1-Ta and HERV-K DNA content and osteocyte-specific gene expression is particularly notable. SOST and MEPE are hallmarks of mature osteocytes and critical regulators of bone remodeling and mineralization both being important in WNT signaling pathway. Reduced expression of these genes is indicative of impaired osteocyte function and is closely linked to skeletal fragility. ^52 53^ The selective correlation of endogenous RT activity with osteocyte markers, but not with early osteoblast genes, suggests that RT-dependent processes are especially relevant for maintaining osteocyte identity and endocrine competence rather than for early lineage commitment.

Using NRTIs as mechanistic tools, we found that RT inhibition selectively disrupted key endocrine functions of mature osteocytes, notably 1,25-dihydroxyvitamin D_3_-induced FGF23 expression and parathyroid hormone–mediated suppression of SOST. These effects occurred with minimal impact on early differentiation markers and only modest effects on mineralization. The results suggest clearly that endogenous RT activity is particularly important for hormone responsiveness and signal integration to maintain the function of mature osteocytes. Among the NRTIs tested, abacavir exerted the strongest inhibitory effects, nearly abolishing vitamin D-dependent FGF23 induction and PTH responsiveness, highlighting differential sensitivities of mammalian RTs to pharmacological inhibition. The NRTIs produced distinct effects in the cultured cells, likely reflecting drug-specific sensitivities of mammalian retroelement RTs. 3TC and TDF have previously been shown to inhibit LINE-1 retrotransposition with different potencies and maximal effects in human HeLa cells.^54^ Moreover, because each NRTI uses different cellular uptake and phosphorylation pathways ^14^ the intracellular levels and stability of their active metabolites may vary in the mouse IDG-SW3 cell line and differ from the effects in human cells.

Several questions remain. The molecular mechanisms by which endogenous RT activity regulates osteocyte function are not yet defined. One possibility is that RT-dependent cDNA intermediates act as regulatory molecules participating to stress-response pathways and thereby modulating osteocyte signaling and adaptation.

Beyond its conceptual implications for bone biology, this work has translational relevance for conditions involving chronic RT inhibition. Long-term exposure to NRTIs is associated with reduced bone mineral density and increased fracture risk, a phenomenon that has remained mechanistically elusive. Our findings provide a plausible biological framework in which inhibition of endogenous, osteocyte-derived RT activity may contribute to impaired endocrine signaling and skeletal fragility. Importantly, the differential effects observed among NRTIs suggest that drug-specific inhibition of mammalian RTs may influence skeletal outcomes, with potential implications for therapeutic decision-making.

From a broader perspective, endogenous RT activity may represent a biomarker of osteocyte function and bone health, as well as a potential therapeutic target. Strategies aimed at preserving or modulating TE-derived RT activity in bone could support osteocyte endocrine function and mitigate bone loss associated with aging or pharmacological RT inhibition.

## Materials and Methods

### Participants and ethics

The iliac bone donors have been described previously in detail ^41^. Briefly, A total of 301 unrelated postmenopausal Norwegian women aged 50-86 years were consecutively recruited at the outpatient clinic of Lovisenberg Deacon Hospital (Oslo). Of these, 178 were excluded due to comorbidities or medications affecting bone metabolism. Among the remaining 123 eligible participants, 23 subsequently declined participation, leaving 100 women who underwent trans-iliac bone biopsies. Of these samples, 84 were suitable for gene expression analyses, while an additional 7 were used for bone histomorphometry. Nine samples were excluded due to RNA degradation.

Based on total hip Dual-Energy X-ray Absorptiometry (DEXA), 18 of 84 women (21.4%) were classified as osteoporotic (T-score ≤ -2.5), 20 as osteopenic (-2.5 < T-score ≤ -1), and 46 had normal or high bone mineral density (T-score > -1.0). Similar distributions were observed at the lumbar spine (L1-L4). Among participants, 21 women with low BMD had experienced at least one low-energy fracture, with the most recent occurring at least two years prior to the study. Previous estrogen users (n = 39) had discontinued therapy for at least two years (with one exception of six months). All donors taking medication or having diseases, other than primary osteoporosis, that are known to affect bone metabolism were excluded. The presence of bone-impairing diseases/conditions was excluded by extensive biochemical serum and urine analyses supported by X-ray examinations. The site-specific BMD of all donors was evaluated using Lunar Prodigy DEXA (GE Lunar, Madison, WI, USA) following the manufacturer’s instructions. The precision of the instrument for measuring the lumbar spine (L_2_-L_4_) and hip BMD was 1.7 and 1.1%, respectively. The study was approved by the Norwegian Regional Ethics Committee (REK no 2010/2539, Norway). All volunteers gave their written informed consent, and sampling and procedures were according to the Act of Biobanking in Norway.

### DNA extraction and quantification

DNA was isolated from the same bone biopsy samples used previously for RNA extraction ^41^. Briefly, liquid nitrogen– cooled biopsies were pulverized in a mortar and homogenized in TRIzol reagent (Cat. No. 15596026; Invitrogen) following the manufacturer’s protocol for RNA isolation. After chloroform extraction, phase separation yielded an interphase containing DNA between the aqueous and organic layers. The DNA was subsequently recovered from this interphase according to the TRIzol reagent manufacturer’s instructions. DNA concentration and purity were determined using a DeNovix microvolume spectrophotometer (DeNovix Inc.). The iliac bone biopsies contained adherent muscle tissue on the visceral side. This muscle tissue was carefully separated from the bone biopsies, and DNA was isolated using the same protocol as applied for bone DNA. In addition, peripheral blood samples were obtained from the donors in the morning after an overnight fast. Genomic DNA was extracted from EDTA-anticoagulated whole blood using the QIAamp DNA Blood Mini Kit (Qiagen, Hilden, Germany) according to the manufacturer’s instructions.

### L1-Ta and HERV-K DNA content quantification

L1-Ta and HERV-K DNA content was quantified using a Bio-Rad CFX96 Real-Time System. In each reaction, 4.5 μl (0.025 ng) of genomic DNA was added to 5 μl of Luna Universal qPCR Master Mix (NEB) and 0.5 μl of primer mix (final concentration 0.4 μM). Satellite DNA (SATA) content was used for normalization. PCR thermal profile was the following: 95 °C for 1’, followed by 45 cycles of 15’’ at 95 °C and 30’’ at 60 °C. Quantification was performed with the ΔΔCt method. Primer sequences can be found in Table S3.

### RNA extraction and cDNA preparation

IDG-SW3 cells were washed three times with 1× PBS and incubated with 40mM EDTA for 5 minutes to dissolve mineral deposits. Cells were then washed three times with 1× PBS harvested and resuspended in 1 ml of QIAzol Lysis reagent (Qiagen, Cat. No. 79306). Total RNA was then purified using the RNeasy Plus Mini kit (Qiagen, cat. No. 74134) with minimal modifications to the manufacturer’s instructions. DNase treatment (RNase-free DNase set, Qiagen, Cat. No. 79254) was performed to remove any residual DNA. RNA quality and concentration were checked using a Nanodrop™ 2000 spectrophotometer (Thermo Fisher). cDNA was synthesized from 200 ng of each RNA sample using a Superscript III first-strand cDNA synthesis system (Thermo Fisher, cat. No. 18080051) according to the manufacturer’s protocol. Quantification was performed with the ΔΔCt method. Primer sequences can be found in Table S3.

### Microarray analysis

Double-stranded cDNA and biotin-labeled cRNA probes were made from 5 μg of total RNA by use of GeneChip® Expression 3′ Amplification One-Cycle Target Labeling Kit (Affymetrix). The cRNA was hybridized to HG-U133 plus 2.0 chips (Affymetrix) followed by washing and staining on the GeneChips Fluidics Station 450 (Affymetrix) according to manufacturer’s instructions. The chips were scanned on the Affymetrix GeneChip Scanner 3000. RNA probe quality assessment and data pre-processing and evaluation have been extensively described in Reppe et al., 2010 ^41^. Data are available at the European Bioinformatics Institute (EMBL-EBI) ArrayExpress repository, ID: E-MEXP-1618.

### IDG-SW3 cell culture

Cells were expanded in permissive conditions (33°C in αMEM with 10% FBS, 100 units/ml penicillin, 50 μg/ml streptomycin, and 50 U/ml IFN-γ (Sigma, Cat. No. IF005)) on rat tail type I collagen-coated plates. To induce osteogenesis, cells were plated at 8,000 cells/cm^2^ in osteogenic conditions (37°C with 50 μg/ml ascorbic acid (Sigma, Cat. No. A4544) and 4 mM β-glycerophosphate (Sigma, Cat. No. G9422) in the absence of IFN-γ). Following osteogenic induction, 3TC (Sigma, Cat. No. L1295) and ABC (Sigma, Cat. No. SML0089) were administered daily at the concentration of 5 μM and 10 μM, respectively; TDF (Sigma, Cat. No. SML1795) was administered every 3 days at the concentration of 10 μM. At day 19 of differentiation, cells were treated with either 10 nM PTH (Merck, Cat. No 05-23-5501) or 10 nM 1,25-dihydroxyvitamin D_3_ (Cambridge Isotope Laboratories, Cat. No. DLM-9107-B) for 48 hours. Dosages were calculated based on the IC50 values.

### Gene expression analysis

Total RNA was extracted from cells using the Monarch Total RNA Miniprep Kit (NEB). An optional DNase treatment was performed. RNA quality and concentration were measured using a Nanodrop spectrophotometer. cDNA was produced using the SuperScript™ III First-Strand Synthesis SuperMix (Invitrogen). Gene expression was quantified using a Bio-Rad CFX96 Real-Time System. In each reaction, 4.5 μl (4.5 ng) of cDNA was added to 5 μl of Luna Universal qPCR Master Mix (NEB) and 0.5 μl of primer mix (final concentration 0.4 μM). PCR thermal profile was the following: 95 °C for 1’, followed by 45 cycles of 15’’ at 95 °C and 30’’ at 60 °C. 18s RNA was used for normalization. Quantification was performed with the ΔΔCt method. Primer sequences can be found in Table S3.

### Alizarin Red staining

Cells were washed with 1× PBS and fixed with 4% (v/v) formaldehyde (Sigma, USA, Cat. No. F8775) in 1× PBS for 15 min. After washing twice with ddH_2_O, 40mM Alizarin Red staining S (ScienCell, Cat. No.8678) was added for 20 min. Afterward, cells were washed four times with ddH_2_O, dried, and stored in the dark until image acquisition. For absorbance measurement, Alizarin Red S staining quantification assay (ScienCell, Cat. No.8678) was used following manufacturer’s instructions. Absorbance was measured at 405 nm with GloMax® discover microplate reader (Promega, USA).

### Statistical analyses

T-test was used in all DNA content and gene expression analyses, and simple linear regression was used in all correlation analyses. Correlation coefficient (r) and P values are shown in each plot. \**P* < 0.05, \*\**P* < 0.005, \*\*\**P* < 0.0005, \*\*\*\**P* < 0.00005.

### Contributors

AM, SR, KMG and VO contributed to the conception and design of the study. KMG and SR contributed to the clinical data collection. AM and SR contributed to biopsies analysis. AM, GM and MD contributed to *in vitro* experiments, validation and data analysis. AM contributed to drafting and writing the manuscript and producing figures and tables. AM, SR, GM and MD had access and verified all the data. All authors were involved in the data interpretation, review, and approval of the manuscript. All authors read and approved the final version of the manuscript.

The authors declare no conflict of interest.

## Acknowledgements

This research was supported by KAUST BAS/1/1037-01-01, the South East Norway Health Authority and Oslo University Hospital, Ullevaal (52009/8029); The 6th EU Framework Program (LSHM-CT-2003-502941); Legat til Forskning, Lovisenberg Diaconal Hospital.

We thank Prof. Lynda F. Bonewald for kindly providing IDG-SW3 cell lines.

