## Supplemental data for "Endogenous retrotransposon activity supports osteocyte function and links antiretroviral therapy to bone loss"

Figure S1

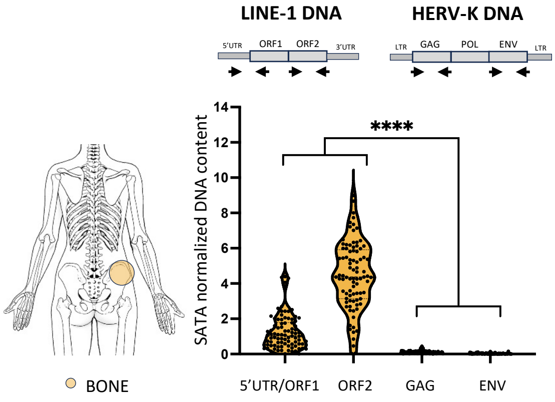


**Fig. S1 Comparison of L1-Ta and HERV-K DNA content in trans-iliac bone biopsies.**
qPCR analysis of L1-Ta and HERV-K DNA content in bone biopsies. Values are plotted over the average of L1-Ta (5'/ORF1 region). Mean (5'UTR/ORF1= 1; ORF2= 3.550; GAG= 0.0848; ENV= 0.02).

Figure S2

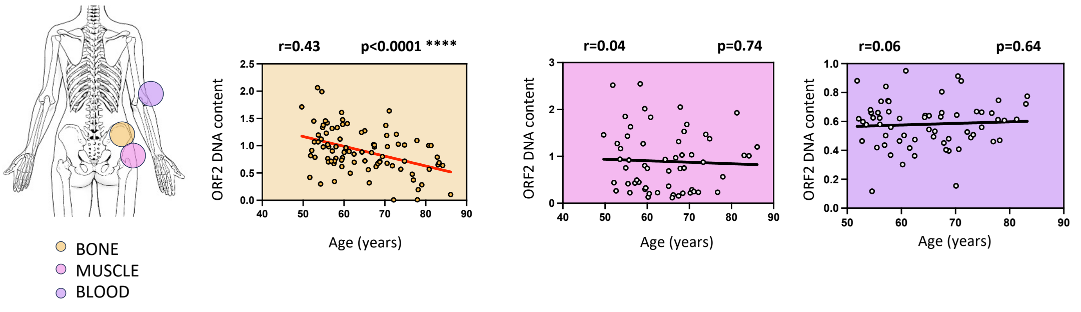


**Fig. S2 Tissue-specific correlation analysis between L1-Ta DNA content and individual age.**
Correlation analysis between L1-Ta DNA content and individual age in bone, muscle and blood biopsies. Correlation coefficient (r) is shown in each plot.

Figure S3

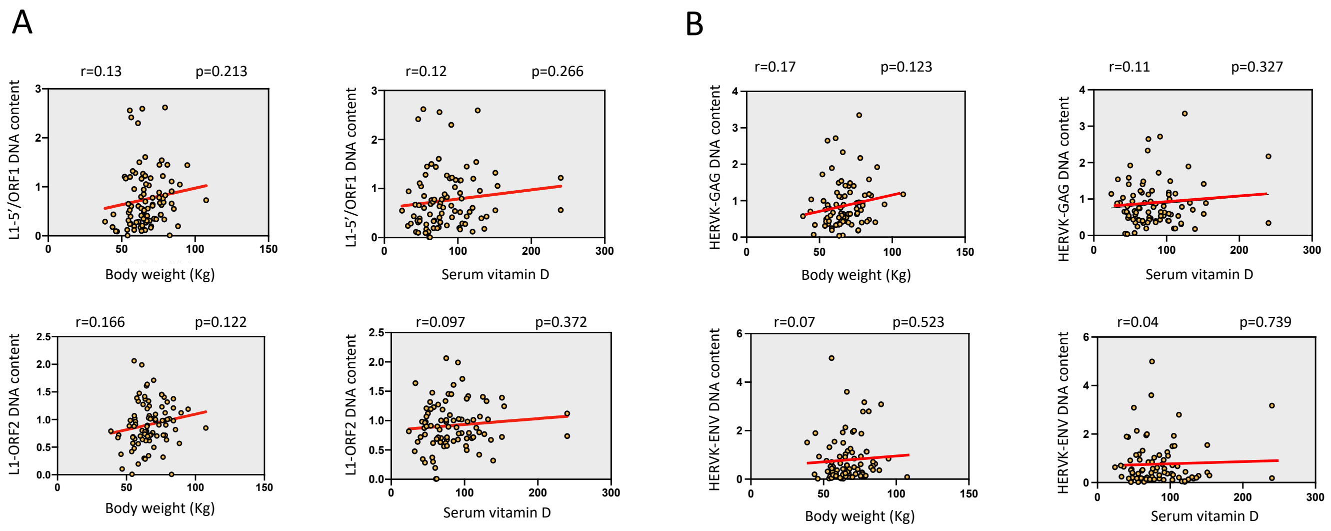


**Fig. S3 Correlation analysis between L1-Ta and HERV-K DNA content and clinical parameters.**
A) Correlation analysis between L1-Ta DNA content, body weight and serum vitamin D. B) Correlation analysis between HERV-K DNA content, body weight and serum vitamin D. Correlation coefficient (r) is shown in each plot.

Figure S4

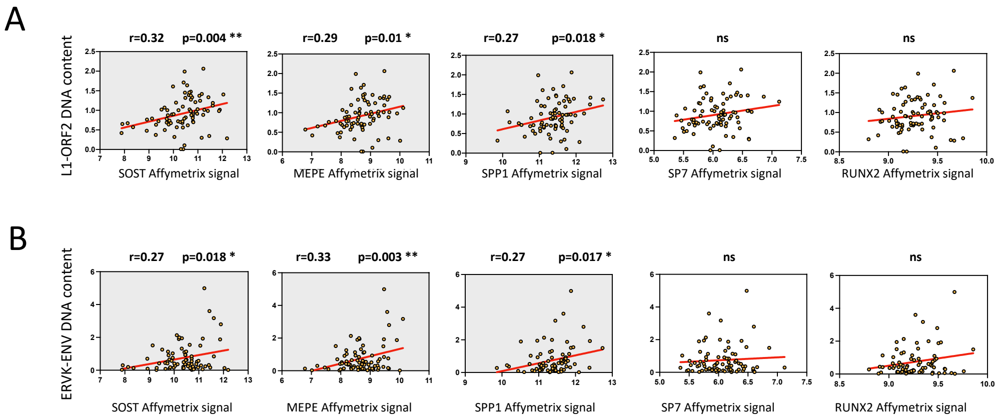


**Fig. S4 Correlation analysis between L1-Ta and HERV-K DNA content and expression of osteocytic genes in trans-iliac bone biopsies.**
A) Correlation analysis between L1-Ta DNA content and the expression of genes downregulated in the biopsies of osteoporotic patients compared to healthy. Correlation coefficient (r) is shown in each plot. B) Correlation analysis between HERV-K DNA content and the expression of genes downregulated in the biopsies of osteoporotic patients compared to healthy. Correlation coefficient (r) is shown in each plot.

**Table S1. Analysis of bone metabolism markers in the serum**

| | Serum osteocalcin (nM) | Serum 1-CTP (mg/l) | Serum bone-spec ALP (u/l) | Urine-Ntx (nM BCE/mM Cr) |
|---|---|---|---|---|
| Laboratory reference ranges | <3.6 | 1.8-5.00 | 11.6-30.6 | <131 |
| Healthy group | 1.333 | 3.553 | 20.59 | 52.28 |
| OP group | 1.693 | 4.683 | 28.95 | 74.54 |
| Difference between means ± SEM | 0.3595 ± 0.1503 | 1.129 ± 0.3580 | 8.364 ± 2.126 | 22.25 ± 8.729 |
| p-value | 0.0097 | 0.0012 | <0.0001 | 0.0066 |

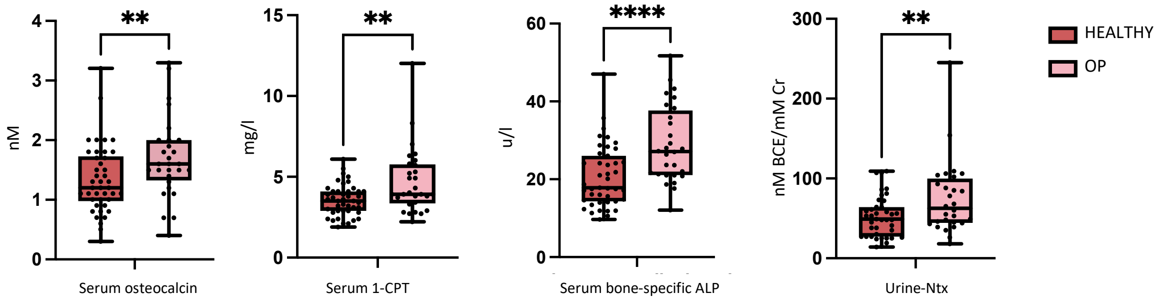

**Table S2. Comparison of cell specific bone related transcript signal levels in trans-iliac bone biopsies between osteoporotic and healthy donors**

| Gene symbol | Gene name | Mean signal level healthy | Mean signal level OP | Difference between means ± SEM | p-value |
|---|---|---|---|---|---|
| | | OSTEOBLAST SPECIFIC TRANSCRIPTS | | | |
| COL1A1 | Collagen Type I Alpha 1 Chain | 12.65 | 12.65 | 0.004379 ± 0.1112 | 0.4844 |
| COL1A2 | Collagen Type I Alpha 2 Chain | 12.73 | 12.7 | 0.02283 ± 0.1606 | 0.4437 |
| BGLAP | Bone Gamma-Carboxyglutamate Protein (osteocalcin) | 11.34 | 11.21 | 0.1272 ± 0.1545 | 0.2067 |
| SP7 | SP7 Transcription Factor (osterix) | 6126 | 5.965 | 0.1609 ± 0.08715 | **0.0348** |
| RUNX2 | RUNX Family Transcription Factor 2 | 9.34 | 9.202 | 0.1378 ± 0.05127 | **0.0046** |
| ALPL | Alkaline Phosphatase, Biomineralization Associated | 5.898 | 5.884 | 0.01426 ± 0.08982 | 0.4372 |
| PTH1R | Parathyroid Hormone 1 Receptor | 6.895 | 6.948 | 0.05231 ± 0.07507 | 0.2443 |
| PTH2R | Parathyroid Hormone 2 Receptor | 5.198 | 5.338 | 0.1404 ± 0.08454 | 0.0508 |
| | | OSTEOCYTE SPECIFIC TRANSCRIPTS | | | |
| SOST | Sclerostin | 10.83 | 9.717 | 1.109 ± 0.1972 | **<0.0001** |
| MEPE | Matrix Extracellular Phosphoglycoprotein | 9.009 | 8.23 | 0.7792 ± 0.1487 | **<0.0001** |
| DMP1 | Dentin Matrix Acidic Phosphoprotein 1 | 6.47 | 6.266 | 0.2043 ± 0.1522 | 0.0922 |
| PTRM | Protein Tyrosine Phosphatase Receptor Type M | 8.07 | 8.054 | 0.01620 ± 0.06915 | 0.4078 |
| | | TRANSCRIPTS IN COMMON BETWEEN OSTEOBLASTS AND OSTEOCYTES | | | |
| SPP1 | Osteopontin | 11.45 | 11.22 | 0.2237 ± 0.1242 | **0.0382** |
| IBSP | Integrin Binding Sialoprotein | 9.85 | 9.621 | 0.2291 ± 0.1593 | 0.0777 |

**Table S3. qPCR primers used in this study**

| Human | | |
|---|---|---|
| Target | Forward sequence | Reverse sequence |
| SATA | GGTCAATGGCAGAAAAGGAAAT | CGCAGTTTGTGGGAATGATTC |
| L1-Ta 5'UTR/ORF1 | GAATGATTTTGACGAGCTGAGAGAA | GTCCTCCCGTAGCTCAGAGTAATT |
| L1-Ta ORF2 | TGCGGAGAAATAGGAACACTTTT | TGAGGAATCGCCACACTGACT |
| HERVK Pol | TTTGCCACTGCCCATTCTCC | AAGACCAATCTGCCATGCAC |
| HERVK Env | TCGAAGCATCAAAAGCCC | GCAGACTAACAACAGACAAAAC |

| Mouse | | |
|---|---|---|
| Target | Forward sequence | Reverse sequence |
| 18s | GTAACCCGTTGAACCCCATT | CCATCCAATCGGTAGTAGCG |
| DMP1 | CACGGACAGCAGTGAATCTGG | GCCGGTCCCCGTACTCTTA |
| MEPE | GGCACCAATGCCGAATGAAG | CCGTGGGATCAGGATACACAG |
| SOST | AGCCTTCAGGAATGATGCCAC | CTTTGGCGTCATAGGGATGGT |
| PHEX | GGGACACTGAAGCCGTACAG | GCCAGCGGAAAGGTGAATG |
| FGF23 | ATGCTAGGGACCTGCCTTAGA | GGAGCCAAGCAATGGGGAA |
